# FitMultiCell: Simulating and parameterizing computational models of multi-scale and multi-cellular processes

**DOI:** 10.1101/2023.02.21.528946

**Authors:** Emad Alamoudi, Yannik Schälte, Robert Müller, Jörn Starruß, Nils Bundgaard, Frederik Graw, Lutz Brusch, Jan Hasenauer

## Abstract

**Motivation:** Biological tissues are dynamic and highly organized. Multi-scale models are helpful tools to analyze and understand the processes determining tissue dynamics. These models usually depend on parameters that need to be inferred from experimental data to achieve a quantitative understanding, to predict the response to perturbations, and to evaluate competing hypotheses. However, even advanced inference approaches such as Approximate Bayesian Computation (ABC) are difficult to apply due to the computational complexity of the simulation of multi-scale models. Thus, there is a need for a scalable pipeline for modeling, simulating, and parameterizing multi-scale models of multi-cellular processes.

**Results:** Here, we present FitMultiCell, a computationally efficient and user-friendly open-source pipeline that can handle the full workflow of modeling, simulating, and parameterizing for multi-scale models of multi-cellular processes. The pipeline is modular and integrates the modeling and simulation tool Morpheus and the statistical inference tool pyABC. The easy integration of high-performance infrastructure allows to scale to computationally expensive problems. The introduction of a novel standard for the formulation of parameter inference problems for multi-scale models additionally ensures reproducibility and reusability. By applying the pipeline to multiple biological problems, we demonstrate its broad applicability, which will benefit in particular image-based systems biology.

**Availability:** FitMultiCell is available open-source at https://gitlab.com/fitmulticell/fit.

**Supplementary information:** Supplementary data are available at https://doi.org/10.5281/zenodo.7646287 online.

## 1 Introduction

Biological tissues are complex entities composed of cells and extracellular components. Tissues occur in different developmental stages and compositions, are highly dynamic and often heavily structured. Specific tissue properties are relevant for a broad range of processes, including tissue homeostasis, viral infection, and tumor development and treatment. For the experimental analysis of biological tissues, imaging techniques are widely used. Common approaches include light and fluorescence microscopy, but more recently also imaging mass cytometry, spatial transcriptomics and related methods are employed (see Lewis et al. [2021] for a review). These experimental techniques provide a variety of quantitative information about biological tissues. Yet, mechanisms underlying specific pattern formation or tissue dynamics often remain elusive. To address these aspects, computational modeling has established itself as a key element to obtain a comprehensive understanding of causal relationships in multi-cellular spatio-temporal systems. Computational models of multi-cellular processes usually capture multiple spatial and temporal scales and describe the emergence of the system’s behavior based on individual building blocks, e.g. individual cells and their interactions. There are several modeling approaches, including discrete, continuous and hybrid model formalisms [Anderson and Quaranta, 2008, Meyer et al., 2020, Waclaw et al., 2015]. Cells and their interactions can, for instance, be described using (energy-based) Cellular Potts Models (CPM) or (force-based) vertex models. Several software packages have been developed, including MCell [Kerr et al., 2008], FLAME [Richmond et al., 2010], CompuCell3D [Swat et al., 2012], Chaste [Mirams et al., 2013], Morpheus [Starruß et al., 2014], and PhysiCell [Ghaffarizadeh et al., 2018] to simplify and standardize the demanding task of model formulation and implementation. Yet, while computational modeling has substantially improved the understanding of multi-cellular systems, it remains a challenging task [Fletcher and Osborne, 2022].

For multi-scale models of multi-cellular processes a key challenge is parameter estimation, the process in which values for the unknown model parameters are determined by fitting the model simulations to experimentally observed data. Parameter estimation is necessary to obtain quantitative models of processes, to analyse processes, to compare competing hypotheses about processes, and to predict the dynamics of processes (e.g., in response to perturbations) [Durso-Cain et al., 2021, Imle et al., 2019, Jagiella et al., 2017, MacLean et al., 2014, Toni et al., 2011]. Systematic, rigorous and uncertainty-aware parameter estimation is only just becoming accessible for multi-scale models with advanced methods and growing computational resources. Reasons for this are that (i) the simulation of multi-scale models accounts for different biophysical processes (e.g., intra- and extracellular signalling as well as physical interaction), which requires computationally efficient simulation algorithms (e.g., hybrid discrete-continuum approaches), and that (ii) the stochasticity of most models (e.g. due to randomness arising from small cell numbers) necessitates repeated simulations.

To cope with the challenges of parameter estimation for advanced computational models, Approximate Bayesian Computation (ABC) methods have been developed and applied [Beaumont et al., 2002, Pritchard et al., 1999, Sisson et al., 2018]. These methods generate samples from an approximation of the Bayesian parameter posterior distribution, without evaluating a likelihood function which can quickly become inaccessible for complex stochastic models. To enable the application of ABC methods to multi-scale models, they have been parallelized on high-performance computing (HPC) infrastructure [Babtie and Stumpf, 2017, Jagiella et al., 2017, Johnston et al., 2014, Sottoriva et al., 2015]. Generic implementations are provided by, in particular, pyABC [Klinger et al., 2018], ABCpy [Dutta et al., 2017], and ELFI [Kangasrääsiö et al., 2016]. Yet, while simulation and inference tools are available, the parameter estimation for multi-scale models of multi-cellular processes still requires a high level of technical expertise. The available tools are not interfaced and code from published application examples is often difficult to reuse and extend. Thus, there is a need for a platform that facilitates and streamlines the entire workflow, from the construction of multi-scale models of various types based on biological principles, to systematic uncertainty-aware data-driven parameter estimation [Hasenauer et al., 2015].

In this work, we introduce the FitMultiCell pipeline, an open-source, user-friendly, and scalable end-to-end platform that integrates modeling, simulation, and parameter estimation, to simplify the analysis of multi-scale and multi-cellular systems. FitMultiCell integrates the state-of-the-art tools Morpheus [Starruß et al., 2014] for model building and simulation, and pyABC [Schälte et al., 2022] for parameter estimation. It builds on an extension of the PEtab standard [Schmiester et al., 2021] for the specification of parameter estimation problems which we additionally introduce in this manuscript. We demonstrate and evaluate the FitMultiCell pipeline using models for tumor growth and liver regeneration.

## 2 Methods

### 2.1 Problem description

We consider the problem of developing quantitative computational models of multi-cellular processes. These models might account for a broad range of biochemical and biophysical processes, including

- cellular signal transduction, metabolism and gene regulation,
- cell movement, proliferation and death, and
- cell-cell communication (e.g., via direct cell-cell-contact areas or the secretion/uptake of biochemical substances).

The spectrum of modeling frameworks for such integrated processes is broad and the models are often stochastic, like the behaviours and decisions by individual biological cells are inherently stochastic. Mathematically, we can write any such model as

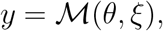

where *y* denotes the vector of observed properties and *θ* denotes the vector of unknown properties of biochemical, biophysical or observation processes (e.g. reaction rates). Process noise – arising, for instance, from low molecular numbers – as well as measurement noise is described via the vector of random variables *ξ*. Marginalizing the model simulation over the random variables *ξ* yields the likelihood *π*(*xy|θ*), which is the conditional probability of observing *y* given *θ*.

Quantitative mathematical modeling requires the inference of the unknown parameters θ from data. In this study, we consider Bayesian inference and aim to approximate the Bayesian posterior distribution

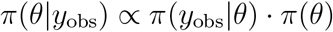

of the unknown model parameters for experimentally observed data *y_obs_* and prior knowledge *π*(*θ*). The posterior distribution encodes all available information about the model parameter and, hence, allows for the assessment of parameter and prediction uncertainties as well as for the design of validation experiments. Experimental data used for the parameterization of the considered models are often obtained using imaging techniques. These can provide information about e.g. spatial profile, temporal changes and changes between conditions. Overall, the spectrum of experimental setups is broad.

### 2.2 FitMultiCell pipeline

To facilitate quantitative computational modeling of multi-cellular processes, we developed the FitMultiCell pipeline (Figure 1). This pipeline supports its users in

1. formulating the modeling problem, i.e.
  a. defining computational models and
  b. defining data obtained from experiments,
2. estimating unknown model parameters, and
3. evaluating parameter and model predictions (including planning validation experiments).

**Figure 1:**
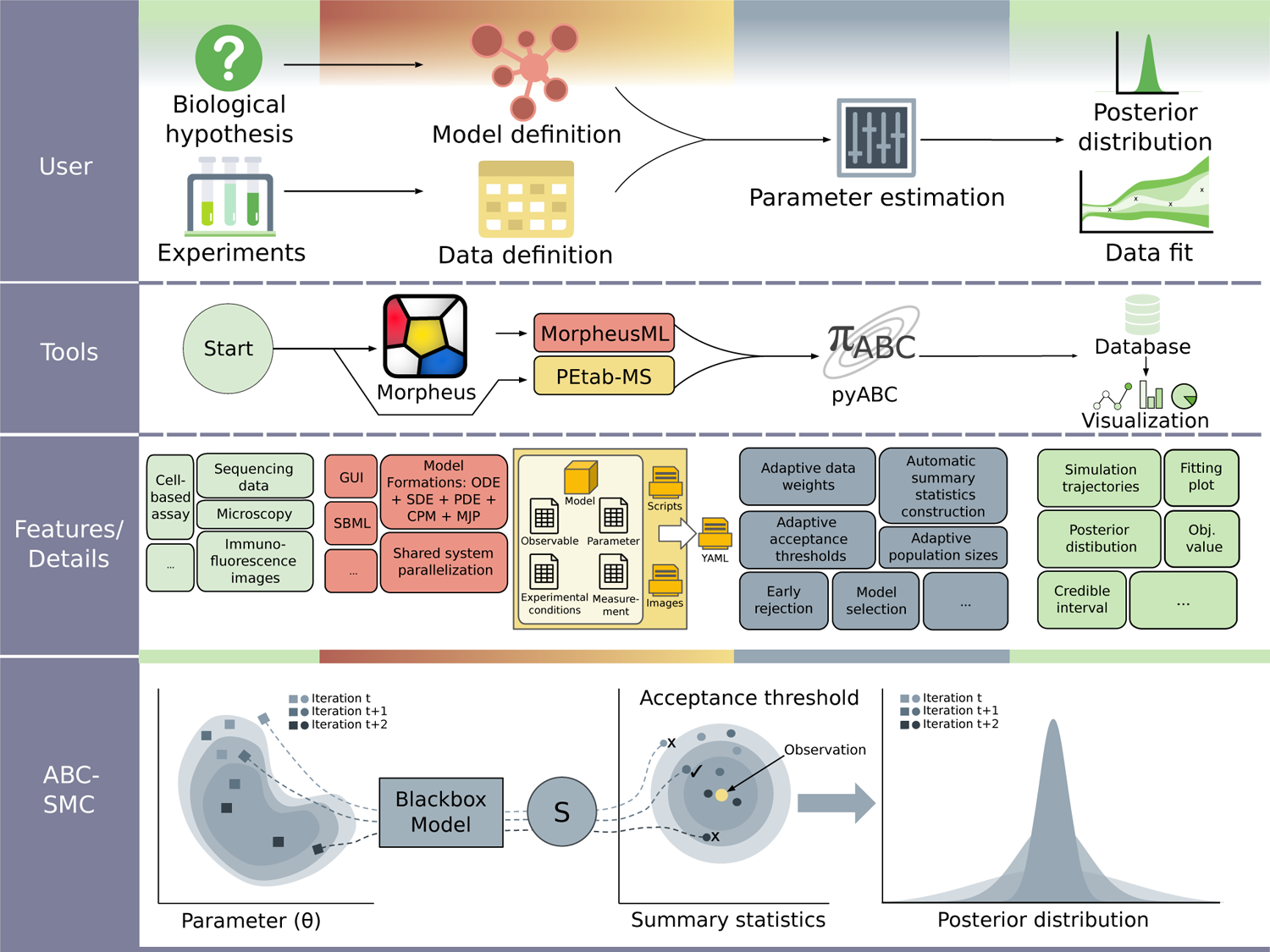
Overview of the FitMultiCell pipeline. First row: User perspective on the biological system, experiment, model formulation, and parameter estimation problem. Second row: Tool and format overview. Third row: Features and details for all components of the workflow. Fourth row: Visualization of the ABC-SMC parameter estimation algorithm.

The FitMultiCell pipeline streamlines the parameter estimation process by providing a tight integration of tools for model specification and simulation, and parameter estimation and reporting. This is achieved using a Python interface with a broad spectrum of functionalities. The current version supports (1) multi-scale modeling and simulation using Morpheus [Starruß et al., 2014], and (2) parameter estimation and uncertainty quantification using pyABC [Klinger et al., 2018]. These state-of-the-art tools cover a broad spectrum of modeling applications; yet, other tools for modeling, simulation and parameter estimation can be easily interfaced.

The FitMultiCell pipeline enables parameter estimation for application problems with multiple experimental conditions, data sets and data types. Furthermore, it offers a broad range of built-in visualization and analysis tools. To facilitate reproducibility and reusability, the FitMultiCell pipeline supports the MorpheusML and the PEtab-MS standards.

In the following, we describe the key components of the FitMultiCell pipeline and their features.

#### 2.2.1 Standardization

The FitMultiCell pipeline uses standardized data formats to ensure interoperability, reproducibility and reusability – important aspects of the FAIR principles [Wilkinson et al., 2016]. The multi-cellular models are encoded using MorpheusML, an established XML-based standard at https://doi.org/10.25504/FAIRsharing.78b6a6. The parameter estimation problems are encoded using the PEtab-MS format, a newly developed tsv-file-based standard.

We developed PEtab-MS as an extension of the Parameter Estimation tabular format (PEtab) [Schmiester et al., 2021], which was developed in the field of ordinary differential equation (ODE) based modeling and is already supported by various simulators and parameter estimation toolboxes. PEtab-MS conserves most core components of PEtab, including the tables defining model parameters and experimental conditions. Yet, PEtab-MS allows for model expression using MorpheusML, flexible data tables (including references to image files), as well as functions for computing summary statistics. These aspects are essential for multi-cellular processes and were not captured by PEtab. The specification of PEtab-MS and a tool for the validation of files can be found at https://gitlab.com/fitmulticell/libpetab-python-MS.

As imaging data are often processed to obtain informative summary statistics, the FitMultiCell pipeline includes a class definition for the construction of summary statistics. Several common summary statistics for imaging data are already provided (see Results section) and customized statistics can be implemented. Furthermore, it is possible to automatically construct informative summary statistics [Fearnhead and Prangle, 2012, Schälte and Hasenauer, 2022].

#### 2.2.2 Simulation

The FitMultiCell pipeline is designed for the simulation-based analysis of multi-cellular processes. Accordingly, we allow for simulation of advanced models M for cells and tissues (using cellular automata and cellular Potts models), extra-cellular concentration fields (using partial differential equations) and intra-cellular dynamics (using ordinary or stochastic differential equations or continuous-time Markov jump processes).

The FitMultiCell pipeline implements a comprehensive interface to Morpheus, including parameter mapping and simulation result extraction. This renders the functionalities of Morpheus (e.g. model construction, simulation and visualization) as well as features such as a comprehensive GUI for rapid prototyping available within the FitMultiCell pipeline. As Morpheus is broadly applicable, widely used for a variety of biological problems, and computationally efficient (see, e.g., Köhn-Luque et al. [2011], Imle et al. [2019] and Vu et al. [2019]), the tight integration of Morpheus in the FitMultiCell pipeline will be beneficial for users.

Additionally, the modular architecture of the FitMultiCell pipeline allows for the interfacing of additional modeling and simulation toolboxes, as well as user-provided simulation codes.

#### 2.2.3 Inference

The FitMultiCell pipeline is designed for simulation-based inference, a class of approaches circumventing the evaluation of the likelihood function. This is important for the study of stochastic multi-cellular processes, but also allows for the application to deterministic models. The FitMultiCell pipeline implements a comprehensive interface to pyABC [Schälte et al., 2022], including parameter and condition mapping for multi-experiment and multi-data type inference. This renders the state-of-the-art Approximate Bayesian Computation Sequential Monte Carlo (ABC-SMC) method easily accessible to users of the FitMultiCell pipeline. ABC-SMC methods approximate the posterior distribution by constructing a sequence of particles which resembles the data (in terms of summary statistics) more and more closely [Toni and Stumpf, 2010]. Well-tested adaptation methods, e.g. for acceptance thresholds, proposal distributions, and population sizes, render pyABC accessible to non-expert users. Furthermore, pyABC allows for exact inference [Schälte and Hasenauer, 2020] and we recently implemented automatic data normalization, summary statistics construction, and measurement noise handling to enable the robust estimation of parameters of multi-scale models [Schälte and Hasenauer, 2022, Schälte et al., 2021]. For the combined use of Morpheus and pyABC, we implemented an early rejection mechanism that discards simulations as early as possible, e.g. if they exceed an upper time limit. This makes the analysis robust to unex-pectedly long-running simulations, e.g. due to an excessive number of cells.

We provided the interface to pyABC as this tool has been successfully used in a broad spectrum of applications, e.g., cardiac electrophysiology [Cantwell et al., 2019], human atrial cells [Houston et al., 2020], cancer [Colom et al., 2021], gene expression [Coulier et al., 2021], universe expansion [Bernardo and Said, 2021], and bee colonies [Minucci et al., 2021]. This underlines its applicability to a broad spectrum of inference tasks for multi-cellular processes. Additionally, the modular architecture of the FitMultiCell pipeline allows for the interfacing of additional inference tools, as well as user-provided inference algorithms.

#### 2.2.4 Distributed execution

Parameter estimation often requires thousands to millions of stochastic model simulations, which is computationally demanding for complex multi-cellular processes. Distributed execution of the computational tasks is thus paramount (Figure 2). Within the FitMultiCell pipeline, parallelization can happen on three levels to efficiently exploit high-performance computing (HPC) infrastructure:

- *Individual simulations* can be parallelized within the employed simulation toolbox. Morpheus supports the use of multiple threads using OpenMP. In addition to ODEs and PDEs, Morpheus also provides a parallel and exact solver for CPMs.
- *Individual summary statistics evaluations* can be parallelized within the FitMultiCell pipeline if multiple individual simulations are required. This is for instance the case if experimental replicates are available for stochastic processes and multiple experimental conditions are considered. As the respective simulations are independent, they can be trivially parallelized across multiple threads.
- *Parameter estimation* can be parallelized within the employed parameter estimation toolbox. PyABC supports single-machine multi-core execution and multi-machine distributed execution. A main process manages the communication across SMC generations and post-processing of accepted particle populations, while the computation-heavy simulations and summary statistic evaluations are delegated to a set of parallel worker processes (see Figure 2). pyABC provides two parallelization strategies, static and dynamic scheduling, which distribute work across computational resources, aiming to minimize the overall CPU time, or the overall wall time, respectively [Klinger et al., 2018].

**Figure 2:**
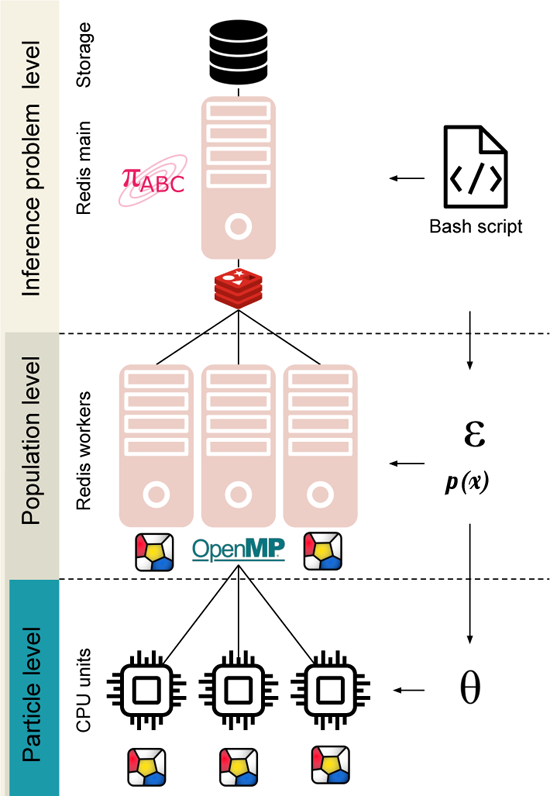
Illustration of the parallel framework of FitMultiCell. ***Inference level:*** A main process manages the pyABC analysis workflow and SQL data storage. ***Population level:*** In each ABC-SMC generation, the main process delegates to a number of distributed worker processes the task of sampling parameters (from proposal distributions *p*(*x*)) and synthetic data sets, until sufficiently many fulfil the generation-specific acceptance criterion (such as the acceptance thresholds 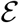). Work distribution, communication, and storage of intermediate results across various computational nodes and processing units are managed via a central Redis server. ***Particle level:*** Each single simulation of synthetic data (or groups of simulations for related perturbation scenarios in a multi-experiment setting) is performed using Morpheus, optionally with shared-memory parallelization via OpenMP.

To facilitate the setup of the required server-worker architecture, the FitMultiCell pipeline comes with bash scripts that handle the resource allocation and communication between nodes, and which can be used to run the pipeline on HPC clusters with minimum code modification. While parallelization is available at multiple levels, the most efficient resource use will often be achieved by dedicating many workers to the particle population and setting Morpheus to single-threaded simulation mode.

#### 2.2.5 Analysis and visualization

The FitMultiCell pipeline provides a GUI for the analysis of the parameter estimation results. This GUI can load the shareable SQL database generated by pyABC, but is flexible enough to visualize the results for other sampling tools after the result files were reformatted. Among other things, the GUI allows for the visualization of (1) samples using uni- and bi-variate plots, (2) sampling diagnostics like the decrease of the acceptance threshold over generations, and (3) the ability to generate the code for any plot that is generated.

### 2.3 Implementation and development

The FitMultiCell pipeline is written in Python and available on GitLab (https://gitlab.com/fitmulticell/fit) and Zenodo (https://doi.org/10.5281/zenodo.7646287) under a BSD-3-Clause license. It is being developed by contributors from three institutions and others are invited to contribute. Code quality is monitored via unit tests and continuous integration. The tool can be installed from the Python Package Index (PyPI). Detailed documentation of the FitMultiCell platform is available at https://fitmulticell.readthedocs.io, which covers installation, setting up the modeling and estimation problem, and running the parameter estimation process, including on HPC and cloud infrastructures.

To ensure ease of use, FitMultiCell pipeline integrates currently only tools which are easy to install and available under permissive licenses:

- Morpheus is available as a git repository (https://gitlab.com/morpheus.lab/morpheus) under the BSD-3-Clause license. It is written in C++ and available for all major operating systems as pre-compiled packages together with training materials and a model repository at (https://morpheus.gitlab.io/).
- pyABC is hosted on GitHub (https://github.com/icb-dcm/pyabc) under the BSD-3-Clause license. It is written in Python and can be installed directly via (PyPI). Its many features are documented at (https://pyabc.readthedocs.io).

## 3 Results

To assess the performance of the FitMultiCell pipeline, we evaluated it using a broad spectrum of test and application examples. In the following, we present its core properties for a few selected models of multi-cellular processes. Several other applications to various multi-cellular problems and different data types have been published separately and are not included here (See Boutillon et al. [2022] Bundgaard et al. [2022]).

All the following results were obtained on the JUWELS standard compute nodes of the Supercomputing Center in Jülich, Germany. Details on the technical specifications are provided in the Supplementary Material.

### 3.1 Standards supported by FitMultiCell pipeline allow for reimplementation of published application problems

The FitMultiCell pipeline allows for the standardized description of models, datasets and parameter estimation problems. To ensure that the supported standards are practically useful, we assessed to which degree we can implement already published application problems. We considered three applications capturing different biological and technical problems: (M1) a model of virus transmission via cell-free virions and cell-to-cell contact [Kumberger et al., 2018]; (M2) a model of tumor spheroid growth [Jagiella et al., 2017]; and (M3) a model describing the mechano-sensing of the metabolic status during liver regeneration [Meyer et al., 2020]. The models M1 and M2 were originally not available in MorpheusML and were reimplemented for the purpose of this study, while M3 was already published in MorpheusML. For all applications, we created the PEtab-MS description of the parameter estimation problems. Details on the application problems are provided below and in the Supplementary Material.

We compared the implementation of the models in MorpheusML with the originally published models and found a good resemblance. Furthermore, PEtab-MS was flexible enough to describe all parameter estimation problems. This suggests that the standards employed in the FitMultiCell pipeline are sufficient for a broad spectrum of applications.

### 3.2 Parallelization in FitMultiCell pipeline provides substantial wall-time reduction

As the efficient use of computing resources is of crucial importance for parameter estimation, we evaluated the parallel efficiency achieved using the FitMultiCell pipeline. We considered a model of the spread of a viral infection within a tissue distinguishing between virus trans-mission via cell-free virions and direct cell-to-cell contact [Kumberger et al., 2018] (Figure 4A and Supplementary Material, Section 4).

**Figure 3:**
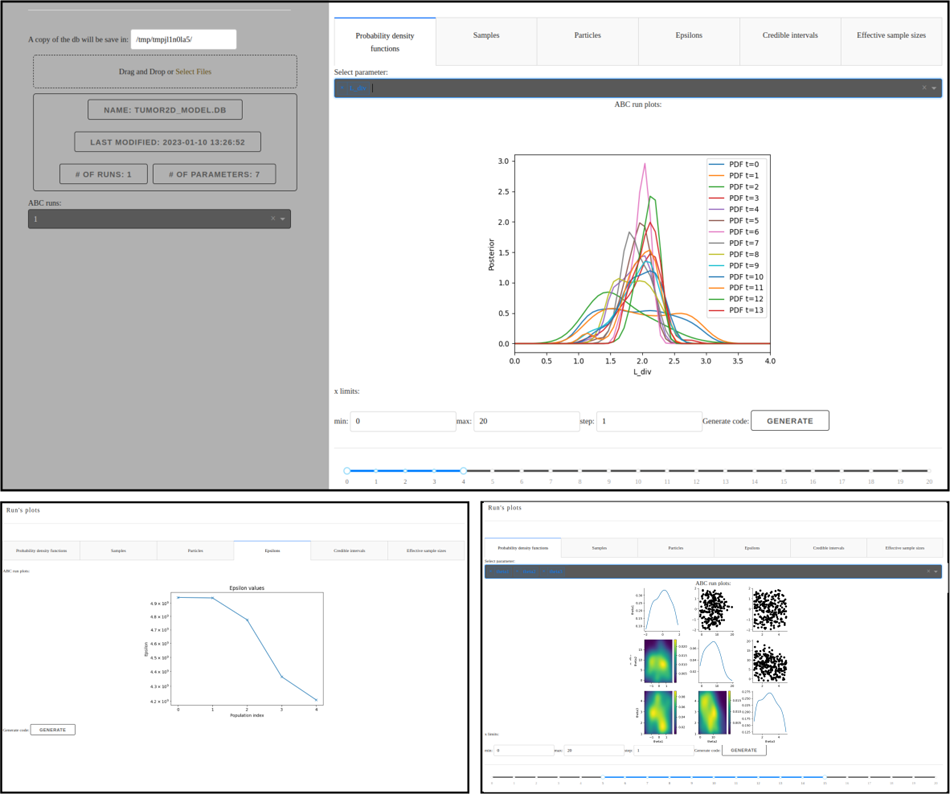
Several screenshots of the visualization and diagnostics GUI.

**Figure 4:**
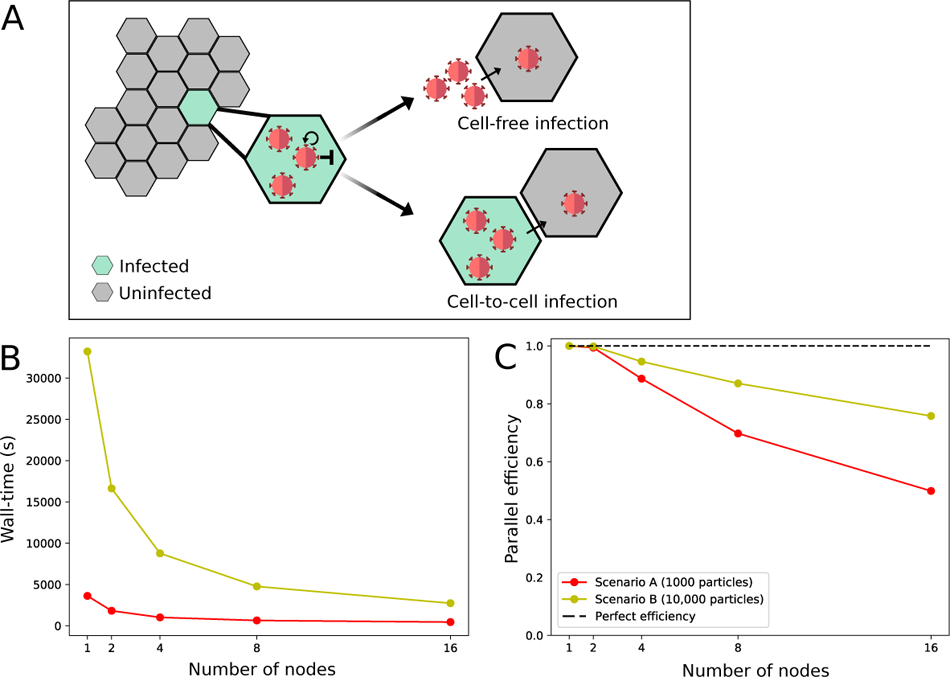
Performance of the parallelized FitMultiCell pipeline. (A) Illustration of the used model M1 for viral transmission within the host tissue. (B) Wall-time of two different fit scenarios across different numbers of used compute nodes, each consisting of 48 CPU cores. (C) Parallel efficiency of the FitMultiCell pipeline. The dashed line indicates perfect efficiency, i.e. a wall-time directly inversely proportional to the number of nodes, compared to single-node execution. For each choice of nodes, three consecutive ABC-SMC generations were run, with population sizes of *N* = 1, 000 (Scenario A) and *N* = 10, 000 (Scenario B).

In the FitMultiCell pipeline, parallelization is available within and across individual simulation particles. For the considered problems, we parallelize exclusively on the population level to provide the most efficient resource usage; however, this may depend on the specific problem.

To assess the parallel efficiency, we performed parameter estimation using the ABC-SMC algorithm implemented in pyABC with the dynamic scheduling option. The algorithms were run for three generations with two different population sizes: (Scenario A) 1,000 particles; and (Scenario B) 10, 000 particles. As higher population sizes improve the posterior approximation but are computationally more demanding, both scenarios are relevant. We performed this task using 1, 2, 4, 8 and 16 compute nodes, each consisting of 48 CPU cores.

For both Scenarios A and B, we observed a substantial reduction in wall-time when using higher numbers of nodes, to a fraction of the single-node execution wall-time which already used 48 CPU cores (Figure 4B). Yet, the parallel efficiency (PE), dropped (Figure 4C) - for the case

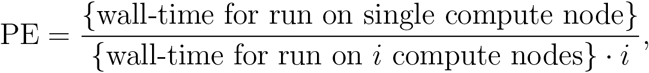

for the case of 16 nodes (768 cores), we found in Scenario A a parallel efficiency of 49% and in Scenario B a parallel efficiency of 75%. The higher parallel efficiency in Scenario B is plausible as individual generations require more time, which reduces the contributions of the tails of generations in which idle times occur due to differences in computation times for individual particles (see also Klinger et al. [2018]).

In summary, our evaluation confirmed an overall good scaling of the FitMultiCell pipeline, yielding a wall-time reduction of several ten-fold compared to a single-node execution and several hundred-fold compared to single-core execution. As the number of proposed particles required to generate a specific number of accepted particles increases over generations, the parallel efficiency for production runs with the usual 15 to 40 generations should be higher than what has been observed here.

### 3.3 FitMultiCell pipeline facilitates the study of heterogeneous datasets via automatically tuned algorithms

To assess the benefit of the tight integration of modeling and parameter estimation tools in the FitMultiCell pipeline, and the availability of advanced inference algorithms, we considered the model of tumor spheroid growth [Jagiella et al., 2017] (Figure 5(left)). The original model was used to study growth control mechanisms and the effect of nutrient supply [Jagiella et al., 2016, 2017]. Yet, the source code of this multi-cellular model entangled with its simulator is highly specific and no longer maintained, complicating the use of this model in further studies.

**Figure 5:**
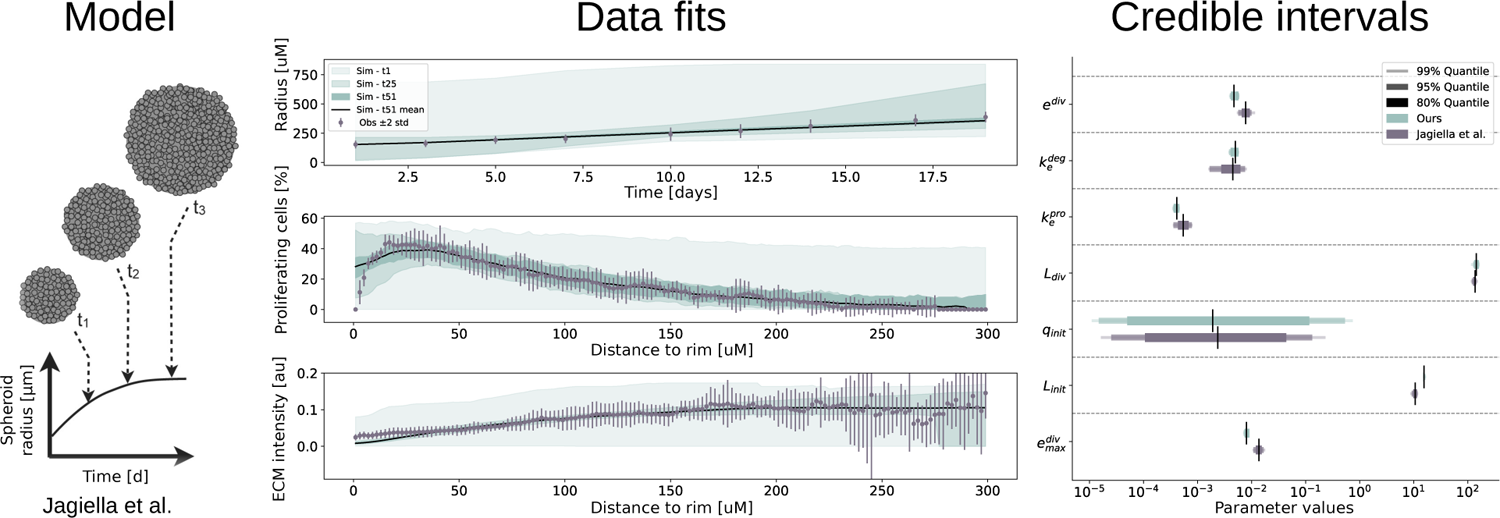
Result of FitMultiCell pipeline for the model of tumor spheroid growth. Illustration of the model (Left). Experimental data (gray) and simulation results (green) (middle). For the simulations associated with different generations the interval between the 2.5th and the 97.5th percentile are shown. Credible interval for 7 estimated parameters obtained using the original implementation [Jagiella et al., 2017] (gray) and the FitMultiCell pipeline (green) (right).

We implemented the model in MorpheusML and made it available through a public model repository (https://identifiers.org/morpheus/M0007). The cell population was described using a stochastic CPM which accounts for cell division and death as well as cell-cell interactions, while the concentrations of extracellular substances were described using partial differential equations (PDEs). The estimation problem for the seven unknown parameters was encoded using PEtab-MS. We considered all available datasets: a time-course for the tumor spheroid radius determined by bright field microscopy; and snapshots for radial profiles of markers for proliferating cells as well as extra-cellular matrix abundance determined from fluorescence microscopy.

In our previous publication considering the parameter estimation problem [Jagiella et al., 2017], we manually defined weights to quantify the relative importance of different summary statistics for the heterogeneous dataset. As this was a time-consuming process, here we employed a fully automatic approach for summary statistics weighting based on inverse regression (see Supplementary Material, Section 2 for details), which is available via the FitMultiCell pipeline. The specification of the parameter estimation task does only require a few lines of Python code given MorpheusML and PEtab-MS files.

We ran the ABC-SMC algorithm with a population size of 500 particles and set as stopping condition a maximum wall-time of 48h, within which the pipeline was able to finish 46 generations. The comparison of simulated and observed summary statistics showed that the model is able to fit the data accurately, with substantially improving fit quality in later ABC-SMC generations (Figure 5 (middle)). Exception are, similar to the original model, the fraction of proliferating cells and the ECM density at small distances to the rim, which the model over- and underestimates, respectively, in correspondence to the original analysis [Jagiella et al., 2017], indicating that the model needs to be refined in this regard. The analysis of credible intervals reveals that with the exception of the initial fraction of quiescent cells c_init_, all parameters are identifiable with small uncertainties. The parameter estimates and uncertainty intervals obtained using our model are in excellent agreement with those in the original analysis (Figure 5(right)). Hence, the FitMultiCell pipeline was able to reproduce the original results, while simplifying implementation and reducing the need for manual tuning.

### 3.4 FitMultiCell pipeline facilitates parameter estimation for new applications

The parameters of multi-scale processes are often still adapted manually, e.g. due to the difficulty of setting up proper parameter estimation and its computational cost. As this involves manual work, it can lead to non-reproducible results and does not provide any information about parameter uncertainties. We assessed whether the FitMultiCell pipeline can easily replace manual parameter tuning. We studied a model describing the mechanosensing of the metabolic status during liver regeneration after partial hepatectomy in mice, simulating the reaction network dynamics in hepatocytes along the central-portal axis of a liver lobule by a spatial array of coupled ODEs with spatially heterogeneous inputs [Meyer et al., 2020]. For this model, the parameters were previously determined manually using data for two observables: the concentration of the YAP protein in the nucleus of hepatocytes (NYAP), and the total concentration of YAP protein in hepatocytes (TYAP), see Figure 6 (left).

**Figure 6:**
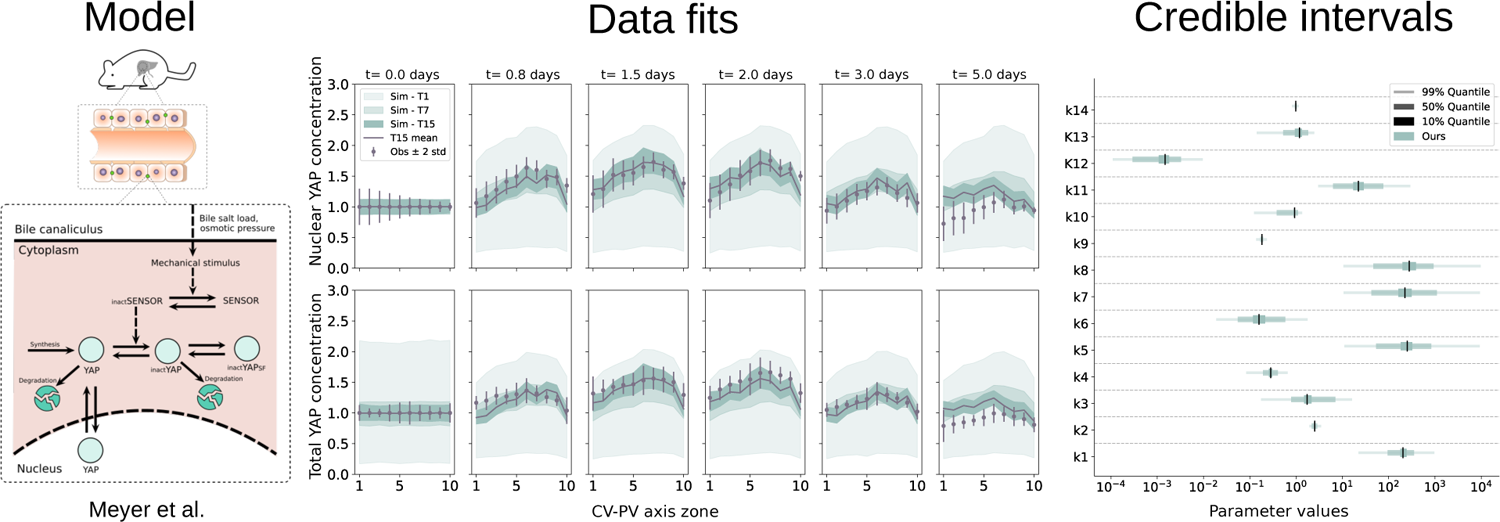
Result of FitMultiCell pipeline for the model of mechano-sensing of the metabolic status during liver regeneration. Illustration of the model (left). Experimental data (gray) and simulation results (green) for different combinations of tissue location (from 1-10) and time point (t0-t5.0 days) (middle). For the simulations associated with different generations the interval between the 2.5th and the 97.5th percentile are shown. Credible interval for 14 estimated parameters (right).

The model was already implemented in MorpheusML and is available from a public model repository (https://identifiers.org/morpheus/M7990). Yet, in exchange with the authors of the original publication, we decided (1) to relax the quasi-steady-state assumption of the intracellular dynamics and performed a full spatio-temporal simulation, (2) to extend the biochemical model by an observer model of the microscopy setup that introduces additional scaling parameters between protein concentration and observed fluorescence intensity, and (3) to account for the measurement noise. Complementary, we created the PEtab-MS files encoding the estimation problem for the 14 unknown parameters. Previously, due to the employed quasi-steady-state assumption, only the ratios of forward and backward rate constants were identifiable.

We ran the ABC-SMC inference with a population size of 1,000 particles and a wall-time limit of 24h. To save computational resources, we employed an early rejection strategy to reject particles based on a maximum runtime of 15min for individual simulations not matching the data. The simulated trajectories in later generations of the inference process fitted the data for TYAP and NYAP mostly well (Figure 6 (middle)). An exception were the pericentral locations (CV-PV zones 1-5) at the last time point *t* = 5.0*days*, where the simulations couldn’t match the low concentrations present in the data, hinting at a transition to homeostasis and a time-dependence of some parameters over longer time scales of many days. The credible intervals indicate that all 14 parameters are identifiable (Figure 6 (right)). Interestingly, the parameter describing the fluorescence intensity normalization (*k*14) was estimated close to one, which supports the hypothesis that both microscopy-based observables can be treated with the same normalization [Meyer et al., 2020]. Moreover, the ratios of rate constants for reversible reactions were found to be on the same order of magnitude as estimated using the quasi-steady state assumption in [Meyer et al., 2020]. Hence, our study underpinned previous conclusions [Meyer et al., 2020], but also demonstrated that the FitMultiCell pipeline allows directly for an uncertainty-aware study of multi-cellular processes.

## 4 Discussion

Quantitative data-based modeling of multi-cellular processes is challenging, because the models are often stochastic and computationally demanding. Furthermore, due to a lack of standards, the reusability of codes and pipelines is limited. Motivated by these issues, we developed the FitMultiCell pipeline, which covers the entire workflow of model development, simulation, and systematic parameter inference (on HPC or cloud infrastructures) based on standardized input formats. Thereby, it contributes to the accelerated testing of biological hypotheses.

We used the FitMultiCell pipeline to study various application problems. This demonstrated that the proposed pipeline is widely applicable and can recover and confirm previous results. Further, we showed that it can replace manual parameter tuning, thus accelerating and solidifying the quantitative modeling of multi-cellular processes. Its modular implementation scales to HPC infrastructures and thereby facilitates inference for computationally expensive problems.

While the FitMultiCell pipeline already provides the necessary features, there are multiple directions in which it can be developed further. (1) Statistical inference for computationally expensive models can become challenging even when using massive parallelization. To address this, the use of cheaper surrogate models could be investigated for guiding parameter search and reducing the number of model simulations [Prangle, 2016, Prescott and Baker, 2021]. Furthermore, one could make use of the idle time at the end of generations, e.g. by presampling the next generation and subsequently correcting for bias. (2) Methods for the automatic construction of summary statistics need to be explored. While it is possible to construct summary statistics semi-automatically [Blum et al., 2013] for the type of image data frequently accompanying models of multi-cellular processes, automatic construction based on machine learning approaches, more specifically convolutional networks with prior information gained from transfer learning, could substantially improve statistics quality at decreased training cost. (3) Extension of the FitMultiCell pipeline to additional modeling, simulation and parameter estimation frameworks will increase its applicability. While Morpheus and pyABC cover already a wide range of applications, an integration of further simulation tools (e.g. Ghaffarizadeh et al. [2018], Merks et al. [2011], Mirams et al. [2013], Swat et al. [2012]) and inference tools (e.g. Dutta et al. [2017], Kangasrääsiö et al. [2016]) could facilitate the use of complementary methods and thus grow the application spectrum further.

In conclusion, we illustrated that standardized workflows for the quantitative modeling of multi-cellular processes are feasible. The FitMultiCell pipeline and the standard PEtab-MS provide starting points for further development. Already in its current form, a broad spectrum of projects can profit from them to achieve systematic and scalable inference of unknown parameters.

## Acknowledgements

We acknowledge fruitful discussions with colleagues at Morpheus.lab and the Gauss Centre for Supercomputing e.V. (www.gauss-centre.eu) for providing computing time on the GCS Supercomputer JUWELS [Jülich Supercomputing Centre, 2019] at Jülich Supercomputing Centre (JSC).

