## Supplementary Material for "FitMultiCell: Simulating and parameterizing computational models of multi-scale and multi-cellular processes"

[*]To whom correspondence should be addressed.

**4 Scaling study** **5**

### 1 Algorithm implementation of ABC-SMC

ABC is a likelihood-free inference method particularly applicable to complex stochastic models, for which likelihood evaluation is computationally prohibitive. This is often the case for multi-scale models [1]. ABC only requires the ability to generate simulated data from the model given input parameters. In a nutshell, ABC calculates a distance between simulated and observed data and accepts corresponding parameters if the distance is below an acceptance threshold. ABC samples from an approximation to the Bayesian posterior distribution, thus providing uncertainty-aware parameter estimates [2]. ABC is frequently combined with an SMC approach, which allows for efficient gradual reduction of the acceptance threshold over a series of particle populations, and which is straightforward to parallelize [3, 4].

There exist various extensions to the core ABC routine, e.g. allowing to auto-tune hyperparameters [5], adapt distances [2] or population sizes [6] to the problem structure, select acceptance thresholds [7, 8], or learn low-dimensional summary statistics [9]. Many such approaches are implemented in pyABC (`https://github.com/icb-dcm/pyabc`), see [4, 10] for details. In particular, such semi-automatic self-tuned and robust approaches make the tool accessible also to users without expert knowledge.

The sampler implementation uses the *Redis* package (`https://redis.io`) as a broker between the main process and workers on a distributed high-performance computing (HPC) architecture.

### 2 Technical specifications

For our test we used Anaconda 3, Python 3.8, with package versions pyABC 0.10.15, Morpheus 2.2.5. Most analyses were performed on the Juelich Supercomputing Center (JSC), Juwels cluster, standard compute nodes, specification of which are 2× Intel Xeon Platinum 8168 CPU, 2× 24 cores, 2.7 GHz 96 ($12 \times 8$) GB DDR4, 2666 MHz. Each node has 48 cores, equaling the number of workers per node used, as all models were single-threaded.

### 3 Model specification

#### 3.1 (M1) HCV model

A model of the spread of hepatitis C virus among cells was used to test the parallel efficiency of the FitMultiCell pipeline. The model was originally developed by [11] and aims to describe viral spread in a spatially-defined environment by accounting for cell-to-cell (CC) and cell-free infection (CF).

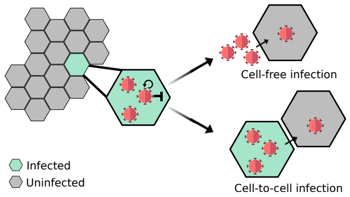

We have re-implemented the model in the FitMultiCell pipeline context. Morpheus was used to generate the MorpheusML model. A smaller version of the model was created, consisting of 721 cells

Table S1: List of fitted parameters of model M1

| Parameter | Description | Prior |
|---|---|---|
| $\log_{10}(sf)$ | Scaling factor of CF infection | U(-3,-1) |
| $\log_{10}(sc)$ | Scaling factor of CC infection | U(-5,0) |

arranged in a hexagonal grid. The model was fitted to synthetic data and the Summary statistics for this model were constructed manually as 1) the number of clusters of infected cells, 2) the number of cells infected by different modes of transmission, and finally 3) the number of cells that can still contribute to CC infection.

##### 3.2 (M2) Tumor Growth Model

The tumor growth model (M2) is our first application to experimental data. The model was developed by [12] and aims to describe the in vitro growth of a tumor spheroid while taking into account its spacial structure.

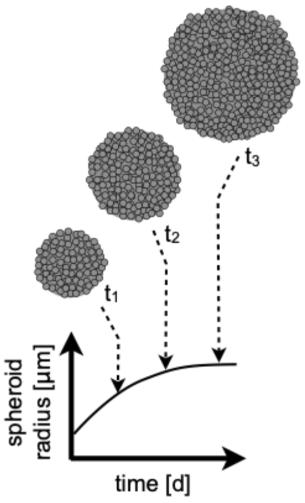

We have re-implemented the model in the FitMultiCell pipeline context. The model was created in Morpheus and encoded as MorpheusML model. The system is built by combining a system of PDE to describe the extracellular matrix and CPM to describe cell configurations and mechanisms like cell division and cell death. For further details about the model see [12], [13].

The model we implemented is a two-dimensional cross-section of the tumor with seven parameters, the details of these parameters are listed in Table S2.

In the early time points, the simulation trajectories of the tumor growth model show stochastic fluctuations, as the number of cells is still small. However, towards the end, the cell numbers are far higher and the fluctuations are much less severe due to the averaging effect of the Law of Large Numbers.

Table S2: List of fitted parameters of model M2. All parameters have been transformed to a logarithmic scale with base 10. The posterior mean and 95% credible intervals are based on the results of the last generation of the analysis.

| Parameter | Description | Prior | Posterior mean | 95% Credible intervals |
|---|---|---|---|---|
| $\log_{10}(k^{\text{div}}_{\text{max}})$ | Division rate | U(-3,-1) | -2.096 | (-2.14, -2.04) |
| $\log_{10}(L_{\text{div}})$ | Division depth | U(-5,0) | 2.15 | (2.10, 2.21) |
| $\log_{10}(L_{\text{init}})$ | Initial spheroid radius | U(1,3) | 1.18 | (1.17, 1.21) |
| $\log_{10}(q_{\text{init}})$ | Initial quiescent cell fraction | U(0,1.2) | -2.72 | (-4.83, -0.27) |
| $\log_{10}(k^{e}_{\text{pro}})$ | ECM production rate | U(-5,0) | -3.38 | (-3.45, -3.33) |
| $\log_{10}(k^{e}_{\text{deg}})$ | ECM degradation rate | U(-5,0) | -2.30 | (-2.41, -2.23) |
| $\log_{10}(e_{\text{div}})$ | ECM division threshold | U(-5,0) | -2.32 | (-2.40, -2.22) |

##### 3.3 (M3) Liver regeneration model

We consider a model of YAP regulation by mechanical stimulation through expansion of the bile canaliculi (BC) [14]. Single realisations of the model vary greatly based on parameter values and need runtimes ranging from 3 to 3,000 seconds. This model has 14 unknown parameters and two observables, namely nuclear YAP and total YAP intensities which were quantified from image tiles covering an entire liver lobule with portal and central veins. The details of the 14 parameters are listed in table S3. The yes-associated protein (YAP) is the downstream effector of the Hippo pathway that plays an important role in liver regeneration and developmental size regulation of many organs.

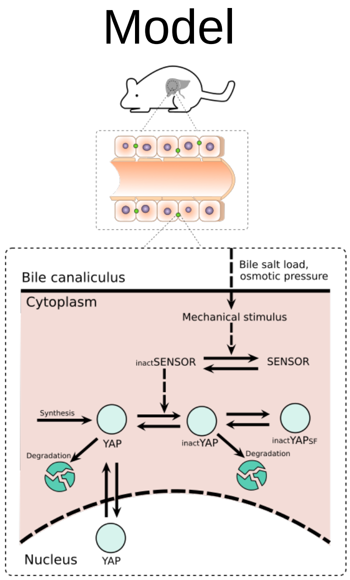

Two sub-models were used to describe the changes in osmotic pressure and the concomitant activation of YAP after partial hepatectomy. The first sub-model is a biophysics-based model to predict the local mechanical stress and apical membrane strain that result from the alteration of osmolyte (bile acid) load in the BC network after partial hepatectomy. It considers the spatial geometry of the BC within the portal and central vein axis of the lobule. Sub-model 2 is a biochemistry-based model that predicts the cellular response of YAP to the local mechanical stress.

To account for measurement noise [15], additive normally distributed noise was added to the simulations, with a standard deviation of $\sigma = 0.0618$ and $\sigma = 0.0501$ for the two observables NYAP and TYAP, respectively, obtained via average standard errors of the mean from the observed data.

Table S3: List of fitted parameters of model M3. All parameters have been transformed to a logarithmic scale with base 10. The posterior mean and 95% credible intervals are based on the results of the last generation of the analysis.

| Parameter | Description | Prior | Posterior mean | 95% Credible intervals |
|---|---|---|---|---|
| $\log_{10}$(k1) | Max. flux of SENSOR activation | U(1,3) | 2.30 | (1.97, 2.54) |
| $\log_{10}$(k2) | Max. flux of SENSOR inactivation | U(0,2) | 0.40 | (0.33, 0.45) |
| $\log_{10}$(k3) | Factor of YAP synthesis rate | U(-0.76,1.23) | 0.23 | (-0.10, 0.84) |
| $\log_{10}$(k4) | YAP inactivation rate | U(-1.95,0.04) | -0.54 | (-0.71, -0.39) |
| $\log_{10}$(k5) | YAP activation rate | U(1,4) | 2.39 | (1.72, 2.91) |
| $\log_{10}$(k6) | Inact. YAP binding rate to SF | U(-1.74,0.25) | -0.80 | (-1.25, -0.23) |
| $\log_{10}$(k7) | Inact. YAP unbin. rate from SF | U(1,4) | 2.34 | (1.62, 3.04) |
| $\log_{10}$(k8) | YAP export rate from nucleus | U(1,4) | 2.44 | (1.66, 2.97) |
| $\log_{10}$(k9) | YAP import rate into nucleus | U(-1.76,0.23) | -0.73 | (-0.77, -0.71) |
| $\log_{10}$(k10) | YAP degradation rate | U(-0.95,1.04) | -0.03 | (-0.41, 0.041) |
| $\log_{10}$(k11) | Inact. YAP degradation rate | U(0.47,2.47) | 1.34 | (0.79, 1.88) |
| $\log_{10}$(k12) | Intensity normalization total | U(-1,1) | -2.82 | (-3.53, -2.48) |
| $\log_{10}$(k13) | M-M const. of SENSOR activation | U(-4,-2) | 0.07 | (-0.27, 0.26) |
| $\log_{10}$(k14) | M-M const. of SENSOR inactivation | U(-1.6,0.40) | -0.005 | (-0.027, 0.015) |

### 4 Scaling study

The details of different scenarios used for the scaling study is presented in table S4. The table contains information regarding the number of populations, population sizes, number of nodes/threads, parallel efficiency, wall time in seconds, and speed-ups, total computation time, the total number of simulations performed, and finally, the acceptance rate.

Efficiency and speed-up are indicated relative to runtimes when using 48 cores as a baseline.

Table S4: Overview of scaling behaviour for different scenarios.

| Scenario | Number of populations | Size of population | Nodes / cores | Parallel efficiency | Wall time (s) | Speed-up | Total computation time (s) | Total simulations | Ave. acceptance rate |
|---|---|---|---|---|---|---|---|---|---|
| A | 3 | 1000 | 1 / 48 | 1 | 3,623.42 | 1 | 173924.16 | 6878 | 0.436 |
| A | 3 | 1000 | 2 / 96 | 0.99 | 1,822.43 | 1.98 | 174953.28 | 6735 | 0.445 |
| A | 3 | 1000 | 4 / 192 | 0.88 | 1,021.42 | 3.54 | 196112.64 | 7290 | 0.411 |
| A | 3 | 1000 | 8 / 384 | 0.69 | 649.23 | 5.58 | 249304.32 | 8016 | 0.374 |
| A | 3 | 1000 | 16 / 768 | 0.49 | 453.91 | 7.98 | 348602.88 | 8884 | 0.337 |
| B | 3 | 10,000 | 1 / 48 | 1 | 33,216 | 1 | 1594368 | 63704 | 0.470 |
| B | 3 | 10,000 | 2 / 96 | 0.99 | 16,637.43 | 1.99 | 1597193.28 | 64203 | 0.467 |
| B | 3 | 10,000 | 4 / 192 | 0.94 | 8,778.62 | 3.78 | 1685495.04 | 64199 | 0.467 |
| B | 3 | 10,000 | 8 / 384 | 0.87 | 4,769.68 | 6.96 | 1831557.12 | 64990 | 0.461 |
| B | 3 | 10,000 | 16 / 768 | 0.75 | 2,738.99 | 12.12 | 2103544.32 | 65826 | 0.455 |

[15] Yannik Schälte and Jan Hasenauer. Efficient exact inference for dynamical systems with noisy measurements using sequential approximate Bayesian computation. *Bioinf.*, 36(Supplement 1):i551–i559, 7 2020.
